# Carpenter ants use diverse antennae sampling strategies to track odor trails

**DOI:** 10.1101/327379

**Authors:** Ryan W. Draft, Matthew R. McGill, Vikrant Kapoor, Venkatesh N. Murthy

**Affiliations:** Center for Brain Science, Harvard University, Cambridge, MA, USA; Department of Molecular & Cellular Biology, Harvard University, Cambridge, MA, USA.; Program in Neuroscience, Harvard Medical School, Boston, MA, USA.

**Keywords:** Ant, Camponotus, Olfaction, Antenna, Trail Tracking, Pheromone

## Abstract

Directed and meaningful animal behavior depends on the ability to sense key features in the environment. Among the different environmental signals, olfactory cues are critically important for foraging, navigation, and social communication in many species, including ants. Ants use their two antennae to explore the olfactory world, but how they do so remains largely unknown. In this study, we use high resolution videography to characterize the antennae dynamics of carpenter ants (*Camponotus pennsylvanicus*). Antennae are highly active during both tracking and exploratory behavior. When tracking, ants used several distinct behavioral strategies with stereotyped antennae sampling patterns (which we call *sinusoidal behavior, probing*, and *trail following*). In all behaviors, left and right antennae movements were anti-correlated, and tracking ants exhibited biases in the use of left vs right antenna to sample the odor trail. These results suggest non-redundant roles for the two antennae. In one of the behavioral modules (*trail following*), ants used both antennae to detect trail edges and direct subsequent turns, suggesting a specialized form of tropotaxis. Lastly, removal of an antenna resulted not only in less accurate tracking but also in changes in the sampling pattern of the remaining antenna. Our quantitative characterization of odor trail tracking lays a foundation to build better models of olfactory sensory processing and sensorimotor behavior in terrestrial insects.

**Summary Statement:** High resolution imaging of antennae reveals distinct patterns of sampling with non-redundant roles in odor tracking.

## Introduction

Ants exhibit remarkably sophisticated and diverse mechanisms of navigation that make use of both internal and external cues (Knaden and Graham, 2016). Sensory signals for navigation come through many modalities, including sight (Narendra, 2017; Graham and Philippides, 2017), touch (Klotz and Reid, 1991; Seidl and Wehner, 2006), and smell (Steck, 2012). Olfaction, in particular, is crucial for the survival and organization of the colony. Ants have highly developed olfactory systems, which they use to communicate with and recognize their colony-mates (Holldobler and Wilson, 1990; Greene and Gordon, 2003) as well as to locate food sources and their nest (Steck, 2011). Ants use species-specific pheromones to create extensive networks of trails that play many roles in colony life, including recruitment of nestmates to active food sources (Traniello, 1977; Holldobler and Wilson, 1990; Morgan, 2009; Czaczkes, 2015). Studies have shown how such trails are generated and adapted over time (Reid, 2010; Czaczkes and Heinze, 2015; Fonio, 2016), and even how they can encode polarity towards or away from the nest site (Jackson, 2004). However, we know very little about how ants are able to accurately track these trails.

Previous studies crossing or amputating antennae have suggested that ants use bilateral odor comparisons (tropotaxis) to follow trails (Hangartner, 1967; Schone, 1984). Theoretical tropotaxis models predict that ants turn towards the antenna with the higher odor concentration while following a trail (Calenbuhr and Deneubourg, 1992). A more recent study looking at ant behavior at a trail bifurcation point has shown that the angle of turning is proportional to the difference in the total quantity of estimated odor on either side of the bifurcation (Perna, 2012). This suggests that ants can discriminate small differences in sensory input to the two antennae and these differences influence motor output. Despite the evidence of bilateral antennae comparisons, little attention has been given to the actual behavioral strategies and the patterns of antennae movements used to actively sample physical space during tracking.

To address this gap in knowledge, we designed a behavioral assay that used high spatial and temporal resolution videography to quantify how ants use their antennae while tracking odor trails. We used the black carpenter ant, *C. pennsylvanicus*, which is widely available, easy to maintain in the laboratory, and which exhibits robust, innate trail tracking behavior. We found that tracking behaviors are much more complex and diverse than previously described. Moreover, antennae usage revealed many interesting properties of insect behavior including stereotyped motor patterns, differences in left-right usage, and behavioral plasticity after amputation. Thus, scent tracking behavior in ants represents a rich model for understanding sensorimotor transformations and for quantitative biology in general.

## Materials and Methods

### Animals

Several colony fragments (~50-100 individuals without queen) were collected in the Middlesex Fells Reservation in Medford, MA during spring and summer months. Individual colony fragments were maintained separately in 17 × 12 × 7.8 inch (Hefty High Rise) plastic containers with sides coated with Fluon (BioQuip, Rancho Dominguez, CA). Containers were lined with leaf-litter and small pieces of dead wood from the collection site. A glass test-tube (6 × 1 inch, VWR) was humidified by adding ~10 mL of water plugged with cotton balls. This tube was placed in the container as a nest area and covered in aluminum foil to block out light. Ants were given fresh water and food (Bhatkar and Whitcomb, 1970) twice per week and kept at constant temperature (23 degrees, 20% humidity) on a 12-hour light-dark cycle (Sylvania Gro-Lux 20 Watt, 24” lights).

### Trail Pheromone

Ants were first anesthetized by placing them in the −20 °C freezer for 5 minutes. Ant gasters were surgically removed with a razor blade. 250 mg of gasters were put into 0.5 mL 70% ethanol and crushed vigorously with a blunt metal rod. The resulting extract was filtered through a 0.45 μm pore filter (VWR sterile syringe filter). Blue fountain pen ink was added (10 μL) for visibility. Ink alone in ethanol did not elicit a tracking response in ants. This liquid was stored at 4 °C and kept up to several months.

### Behavioral Arena and Data Acquisition

A circular platform (1 meter diameter) with 150 mm plastic walls (coated with Fluon) was used as an arena for imaging trail tracking behavior. An enclosure measuring 1 m^3^ was built around the platform to block light and air currents. Three IR light sources (Phenas, 48 LED array 850 nm) were used to illuminate the imaging area. A camera was mounted at the top of the box (DMK 23UM021, 1280×960 pixels, The Imaging Source) to image an 28 × 21 cm area at 0.22 mm/pixel. The bottom of the arena was lined with fresh butcher block paper (Boardwalk, 36-inch × 72 ft) before each imaging session. A piece of letter sized index paper (Limited Papers, 110 lb stock) was taped onto the butcher paper below the field of view of the camera. On this paper, a light pencil trace of the trail (15 cm long) was drawn. Next the trail pheromone was laid down by evenly pipetting the trail extract along the pre-drawn pencil trace with a 10 μL pipet.

An ant was captured at random from a colony and put into a small acrylic box (15 × 13 × 6 cm). This box was then placed in the imaging arena on top of the stock paper. The box had a small opening (2 × 0.5 cm) on one side, which was positioned at the start of the trail. As the ant explored the box, it would find the opening and encounter the trail (this would take 1-2 minutes). This step was necessary to calm and direct the ant, so that it did not immediately run away from the release site and fail to encounter the start of the trail.

Images were acquired at 60 frames per second using IC Capture 2.3 (The Imaging Source) and videos were processed using ImageJ (Schneider, 2012). Videos were down sampled to 30 frames per second for analysis. We stopped recording after the ant tracked the entire trail or abandoned the trail completely. Each ant was given one trial and then returned to the colony. Each trail was used to run multiple ants for up to one hour.

### Data Analysis

All analysis was carried out in MATLAB (Mathworks, Inc.).

### Extracting the Trail Coordinates

To extract trail coordinates, pictures of the trails were taken under visible light before any ants were introduced to the trails. From this image, pixels were classified as part of the trail using an adaptive threshold function in MATLAB. A uniform odor concentration was assumed across all pixels classified as part of the trail.

### Ant Parameters

For each video, a custom-made MATLAB script was used to quantify specific parameters describing the ant in each frame, including the center of mass, the center of the head, the tips of the left and right antennae (absolute and relative to the center of the head), and the body angle of the ant (<u>Movie 1; Movie 2</u>). From these variables, speed, turning angle, and trail overlap of each antenna was derived.

### Antennae Position Distributions

Heat maps were generated by plotting antennae tip position relative to the head position in each frame. To pool data across differently-sized ants, data were normalized by ‘antennae length’ in both X and Y. Antennae length in each dimension was taken to be the farthest X and Y distances, respectively, from the head to the antenna tip over the entire run for each ant. Normalized relative position was displayed as a two-dimensional distribution over 50 bins in both X and Y.

### Behavioral Module Categorization

From the tracking videos, three distinct patterns of behavior (*Probing*, *Sinusoidal, and Trail Following*) were identified. An algorithm was created that used speed and the distance from the head to the trail to segment videos into these different behaviors (Fig. S1). First, if an ant’s speed was less than or equal to 2 pixels/frame (13.2 mm s^−1^) for five consecutive frames, the ant was said to be stationary and these times were marked as Probing. Second, Sinusoidal movement was identified by examining the distance of an ant’s head to the trail over time. To ensure this distance scales with body size, which varies among ants, a relative measure was used as a base unit: a ‘segment’ is the distance from the center of the ant’s head to its centroid (~4.0 mm). The presence of any prominent and distinct peaks in the distance between the ant head and the trail over time were noted (*i.e.*, peaks greater than one-sixth of a ‘segment’ and not closer than two-thirds of a ‘segment’ to each other). To characterize the behavior as Sinusoidal, there had to be at least two marked peaks within 1 second of each other (i.e., periodic movement about the trail). Third, Trail Following was identified as times when the distance from the ant’s head to the trail was less than one-third of a ‘segment’ for at least five frames. Finally, Off Trail behavior was identified as times when the distance from an ant’s head to the trail was greater than two-thirds of a ‘segment’, and any unmarked frames were characterized as Other behavior.

### Trail Overlap and Estimated Odor

A rough estimate of the odor concentration sensed by the antenna was obtained by assuming 1) the antennae are held close to the surface, 2) the antenna’s ability to detect trail odor is approximately a step function of distance, 3) the trail pheromone concentration is uniform among pixels labeled as being part of the trail, and 4) the majority of odorant receptors are at the tip of the antenna (Nakanishi, 2009). A circle of a 10.5 pixel diameter centered at the antenna tip was drawn. The number of trail pixels within this circle is the measure of ‘trail overlap’ or estimated odor concentration (<u>Movie 3</u>). To explore how the width of the step function (the circle) affects our estimation, this trail overlap was plotted as a function of diameter of the circle. Empirically, a circle of diameter 10.5 pixels (2.3 mm) preserved the most information without saturation or redundancy (Fig. S2) and was used for all analyses. For reference, the trail width is approximately 10 pixels (2.0 mm).

### Direction of Antenna Movement

A vector was calculated for the movement of the antenna tip between frames. To prevent alignment imperfections from contributing to the data, data points were discarded if the Euclidean distance of the movement between frames was less than four pixels (0.88 mm). A histogram of the movement angles for each antenna was plotted. The angles from the right antenna were flipped horizontally to match the perspective of the left antennae so data from each behavioral module could be pooled into a single plot.

### Angle of Antenna Joint

To quantify the angle between the flagellar segment and the scape of the antenna, the position of the bend between these two segments was marked manually for both tracking and freely exploring ants. Two lines were drawn between the three marker points: 1) between the antenna tip and the joint and 2) between the joint and the head point. The angle between the two lines was calculated for each frame for both antennae.

### Odor Peak Heights and Widths

To quantify the how the antennae interact with the trail during Probing and Trail Following behavior, the duration an antenna tip stayed on the trail once it made contact was measured as well as the maximum trail overlap it had during contact. This was done two different ways; both gave similar results. In the first method, the ‘findpeaks’ function in MATLAB was used on the trail overlap data. Any peaks closer than 10 frames to another was discarded, and the width of each peak was measured at the half height of the peak. This method removes data from nearby peaks and therefore may underrepresent the duration of continuous encounters with the trail. In the second method, a modified ‘findpeaks’ function was used. The extents of all peaks at the half-height was recorded. Peaks with overlapping extents were considered a single peak and highest peak value was taken as the true peak height. The longest extent was taken as the true peak width. This method was used for data shown in Fig. 5.

### Linear Regression

For Trail Following, a linear regression was performed on the change in an ant’s body angle using the distance of each antenna tip from the trail as predictors. Distance was used instead of our estimated odor measurement because distance is a more continuous variable. Data from frames where both antennae tips were far from the trail (*i.e.*, both had 0 values of trail overlap) was removed. The body angle change at different time points was regressed with respect to antennae tip position (0-8 frames ahead in time; data not shown) to identify any lag between behavioral output and odor input. The highest coefficient of determination (adjusted R^2^) value was found by pairing body angle data from 4 frames after tip position data. In no case did ant speed improve the model as a predictor of body angle change, so it was not included it in our final regression.

### Antennae Correlation

The distances from the left and right antennae tips to the trail (or to the midline) was correlated during ‘bouts’, or sequences of frames for a single behavior of at least 30 frames (1 second). To determine significance, correlation values were first converted using Fisher’s Z-Transformation.

### Left-Right Antenna Bias

To model the expected difference in the left and right antennae trail overlap, two methods were used. In the first, the trail overlap values (right and left) for a given ant were pooled. Then half of these values were randomly assigned to one antenna and the remainder to the other antenna, and then a bias measure was calculated: (sum of left values - sum of right values) / sum of all values. This was repeated 100,000 times to generate a bootstrap distribution. Only ants with more than 200 frames of data were included in the analysis.

The second method took into account the total number of peaks in the trail overlap data, which was obtained using the modified ‘findpeaks’ function described above (see *Odor Peak Heights and Widths)*. The total trail overlap values (right and left) for a given ant were summed and this number was divided evenly into the number of peaks present in the observed data. Each peak was then randomly assigned to either the left or right antenna and a bias measure was calculated as described above. This was repeated 100,000 times to generate a bootstrap distribution. Only ants with more than 200 frames of data were included in the analysis.

### Antenna Removal

For antenna removal experiments, ants were captured from a colony and anesthetized by placing them on a bed of ice (covered by a wet paper towel) until they were immobile. Under a stereomicroscope, one antenna was clipped off at the base of the head using sterile scissors. These ants were then marked for easy identification by applying a spot of nail polish to the abdomen with a toothpick. These ants were tested in the behavioral arena between 2 and 72 hours after surgery.

## Results

In this study, we focused our analysis primarily on straight line trails where all sensory information apart from olfactory cues from the trail (visual and tactile cues) were minimized.

### Antennae sample a wider area of space during trail tracking

To study the differences in antennae usage during trail tracking, we compared the relative antennae positions for ants introduced to an odorant trail (‘Trail Tracking’;<u>Movie 1</u>) with those of ants not given a trail and allowed to run freely in the behavioral arena (‘Exploratory’). We focused on the antennae tips (Fig. 1A), because this is the region of the antennae where odorant receptors are most highly concentrated (Nakanishi, 2009). When tracking odor cues, ants use their antennae to sample a large, arc-like region of space around the midline (where the trail is typically located) (Fig. 1B; <u>Movie 3</u>). During Exploratory behavior, the antennae tips are held within a smaller area located away from the ant’s midline. Although the mean speed of the ant centroid is significantly lower during Trail Tracking relative to the Exploratory behavior (Fig. 1C left, Wilcoxon Rank-Sum, p<0.001), the mean speed of antennae tip movement (relative to the head) is statistically equivalent (Fig. 1C right, Wilcoxon Rank-Sum, p=0.323). Therefore, small antennae movements occur with similar speeds in both conditions, which shows that ants move their antennae actively whether or not they are tracking an odor trail. However, while tracking, the small movements range over a much wider space including the area directly in front of the ant.

**Figure 1.**
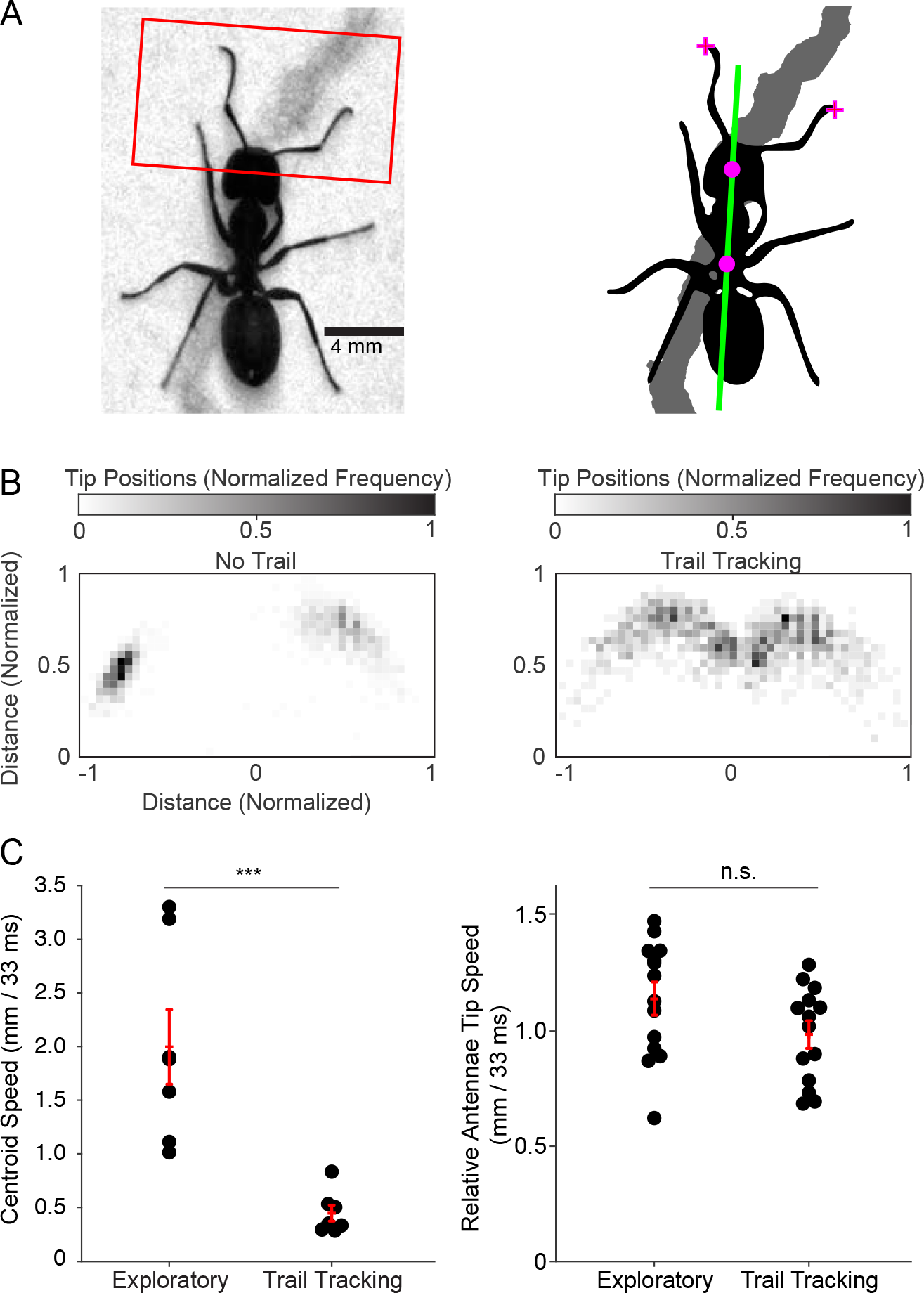
Antennae sample a larger area during trail tracking behavior. A: *(Left)* High resolution image of an ant tracking an odor trail. The red box roughly shows the area over which the antennae tip positions were quantified. *(Right)* A schematic of the same ant showing key parameters used for analysis: antennae tip positions (crosses), head and ant centroids (dots), body axis (green line), and trail pixels above threshold (grey). B: Normalized antennae tip positions from a single ant (1262 data points) without an odor trail (left, Exploratory behavior) versus a single ant (1262 data points) following an odor trail (right, Trail Tracking). C: A comparison of the mean instantaneous speed of the centroid (left) and antennae tips relative to the head (right) during Exploratory behavior and Trail Tracking (Exploratory: n=7 ants; Trail Tracking: n=7 ants). Trail Tracking ants are significantly slower (Wilcoxon Rank-Sum, two-tailed, p<0.001), but no significant difference was found in relative antenna tip speed between the two conditions (Wilcoxon Rank-Sum, two-tailed, p<0.323). Mean and SEM for each distribution is shown in red. *p < 0.05; **p < 0.01; ***p < 0.001.

### Ants exhibit several distinct behavioral strategies during trail tracking

While tracking, ants exhibit substantial differences in their body movements and apparent behavioral states (Fig. 2). Ants moved in sinusoidal paths centered near the trail (Sinusoidal behavior), moved at very slow speeds along the trail (Probing behavior), or followed the trail very closely and accurately (Trail Following). All ant trials included one or more of these three behavioral modules (Fig. S3), which were codified using empirically-derived parameters (see Methods). We marked when ants were far from the trail (Off Trail) and when ants exhibited behaviors that did not conform to the above categorizations (Other behavior). The presence of these easily observable and classifiable behavioral states among a population of ants demonstrates that trail tracking involves multiple strategies and is not a homogeneous behavior.

**Figure 2.**
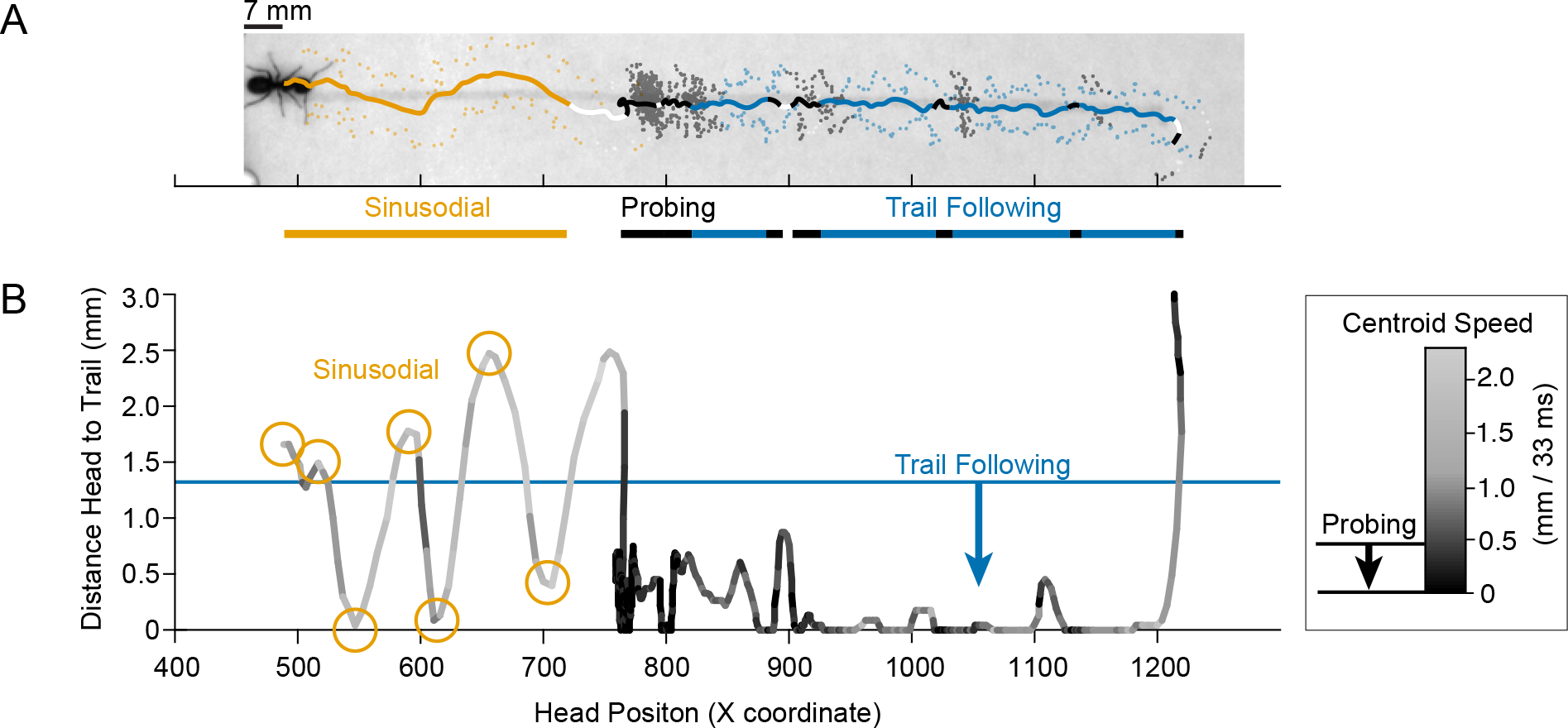
Trail Tracking behavior was separated into three behavioral modules. A: Antennae tip and head positions plotted on an image of the ant and the odor trail. Points are colored according to the behavioral module of the ant in each frame: Light orange: Sinusoidal; Black: Probing; Light blue: Trail Following; White: Other. Bars below the plot show consolidated blocks of the three tracking behavioral modules (Sinusoidal, Probing, and Trail Following). B: For the ant above, a plot of the distance of the head to the trail versus the X coordinate of the image. The curve is pseudo-colored to show the speed of the ant (5-point moving average). Speeds less than 2 pixels/frame (0.44 mm / 33 ms) were used to identify ‘Probing’ behavior. Clusters of large peaks in the distance values were used to identify ‘Sinusoidal’ behavior; distances of the head to the trail less than ~1.3 mm were used to identify ‘Trail Following’ behavior; distances larger than ~2.6 mm were used to identify ‘Off Trail’ behavior. All other frames were marked as ‘Other’. See Methods for details.

### Behavioral modules show differences in antennae sampling and antennae movement patterns

We examined the antennae tip positions (relative to the head) separately for each behavioral module and found robust differences in sampling space (Fig. 3A). Antennae tips are close to each other and close to the head during Probing behavior. In contrast, during Exploratory and Sinusoidal behaviors, the antennae were both farther away from the head and each other. In Trail Following, the antennae spacing is intermediate relative to other behaviors, and antennae are largely excluded from an area at the midline approximately the width of the trail (~2.0 mm). To quantify these differences, we examined the distance of an antenna tip to the head (rho) and the angle of the antenna relative to the ant’s body axis (theta) (Fig. 3B). Antennae tips were closer to the head and had more acute angles relative to the body axis for Probing behavior than for other behavioral modules (Fig. 3C). Trail Following had intermediate values of these parameters, and Sinusoidal and Exploratory behaviors had the largest values. For both rho and theta values, all differences were significant except for the comparison between Sinusoidal and Exploratory behaviors (Kolmogorov-Smirnov, Bonferroni corrected, p<0.0001 and p>0.06, respectively).

**Figure 3.**
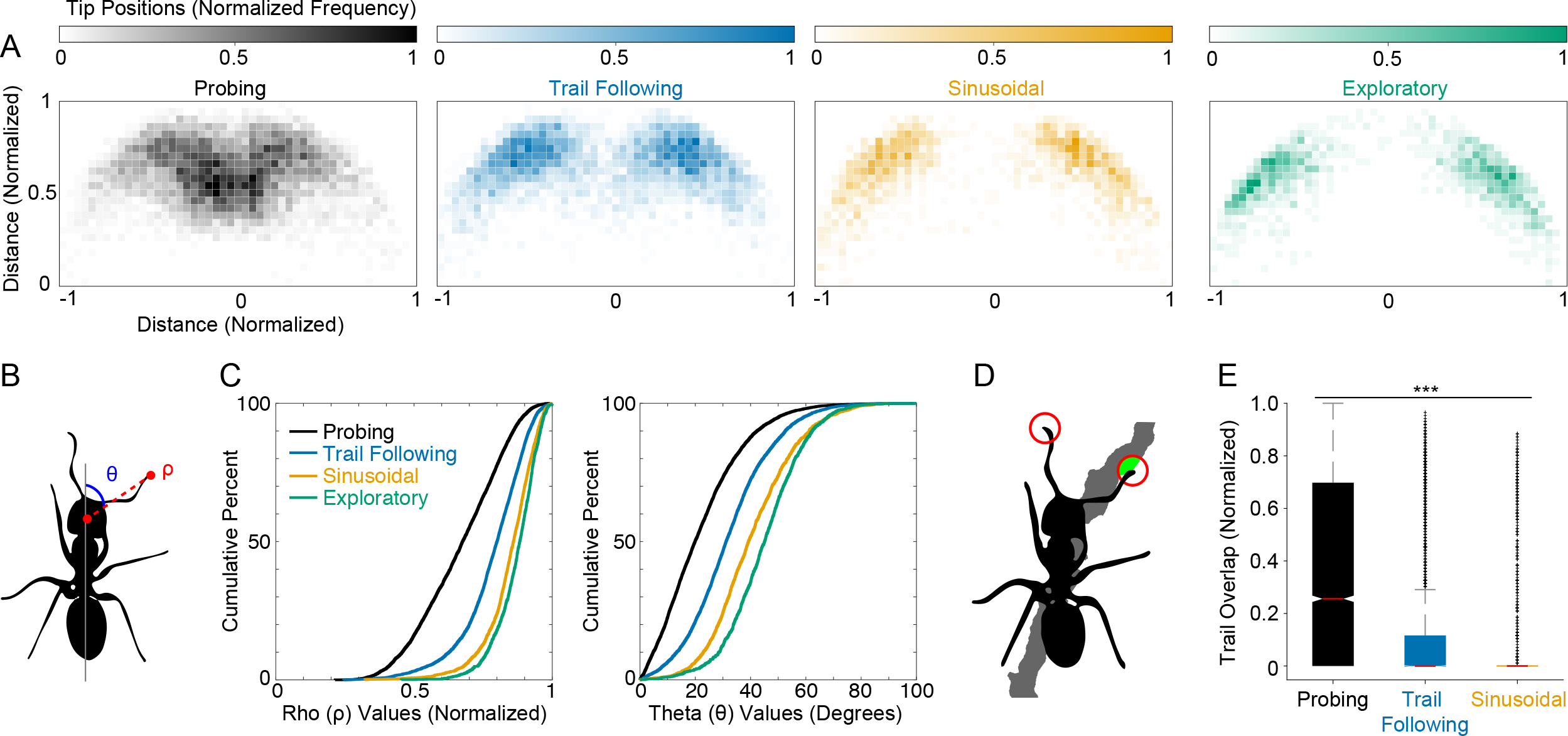
Antennae sample space and receive odor information differently in distinct behavioral modules. A: Antennae tip positions were normalized to the length of each ant’s antennae (1*full antenna length away). Normalized antennae tip positions from Exploratory behavior (n*7 ants, 1420 data points) and Trail Tracking (n*29 ants) separated into behavioral modules (Probing: 8158 data points, Trail Following: 4618 data points, and Sinusoidal: 1822 data points). B. Schematic showing two variables used to quantify the distributions in part A. Rho (ρ) is the distance of the antenna to the head, and theta (θ) is the angle between the body axis and a line connecting the head and the antenna tip. C: Cumulative percent plots of ρ and θ for each behavioral module. All behaviors show distinct distributions except Sinusoidal and Exploratory (Kolmogorov-Smirnov, two-tailed, p < 0.0001 and p > 0.06 respectively, Bonferroni corrected, alpha= 0.008). D: Trail overlap (green) was quantified as the overlap between a 10.5-pixel diameter circle centered around the antenna tip and the trail pixels - see Methods for details. Values were normalized to the maximum possible overlap (89 pixels). E: Trail overlap during the three trail tracking behavioral modules (n=29 ants). Overlap was highest during Probing, followed by Trail Following, then Sinusoidal behavior (Wilcoxon Rank-Sum, two-tailed, p < 0.0001, Bonferroni corrected, alpha= 0.008). *p < 0.05; **p < 0.01; ***p < 0.001.

There is currently no method to measure the actual concentrations of specific chemicals sampled by the antennae. Therefore, to relate the differences in the sampling space to odor sensation, we measured the extent of overlap between a 10.5 pixel diameter circle centered around an antenna tip with the trail (Fig. 3D). Even though the ant was very close to the trail in both Probing and Trail Following behaviors, sampling during Probing resulted in greater antennae overlap with the trail (Fig. 3E). Antennae overlap with the trail was greater for both Probing and Trail Following compared to Sinusoidal behavior, where the ant is often farther than an antenna length from the trail (Fig. 3E, Wilcoxon Rank-Sum, Bonferroni corrected, p<0.0001). This shows that differences in sampling among behavioral modules are likely to result in differences in odor sensation.

To better understand the antennae movements that underlie these sampling differences, we investigated the change in position of antennae tips between subsequent frames (Fig. 4A). These instantaneous movement vectors exhibited three different distributions (Fig. 4B, Kolmogorov-Smirnov, Bonferroni corrected, all pairs p<0.0001 except Sinusoidal and Exploratory: p=0.224). During Probing, ants swept their antennae along a roughly 45 degree-diagonal towards and away from the head. In Exploratory and Sinusoidal behaviors, antennae movements occurred along a roughly 120 degree-diagonal that is approximately perpendicular to the movement patterns during Probing. During Trail Following, the antennae move in a direction perpendicular to the ant’s body axis. Investigation of the sequence of antennae movement vectors confirms that the antennae tips move back and forth between the two peaks shown in the polar histograms across the different behavioral modules (Fig. S4).

**Figure 4.**
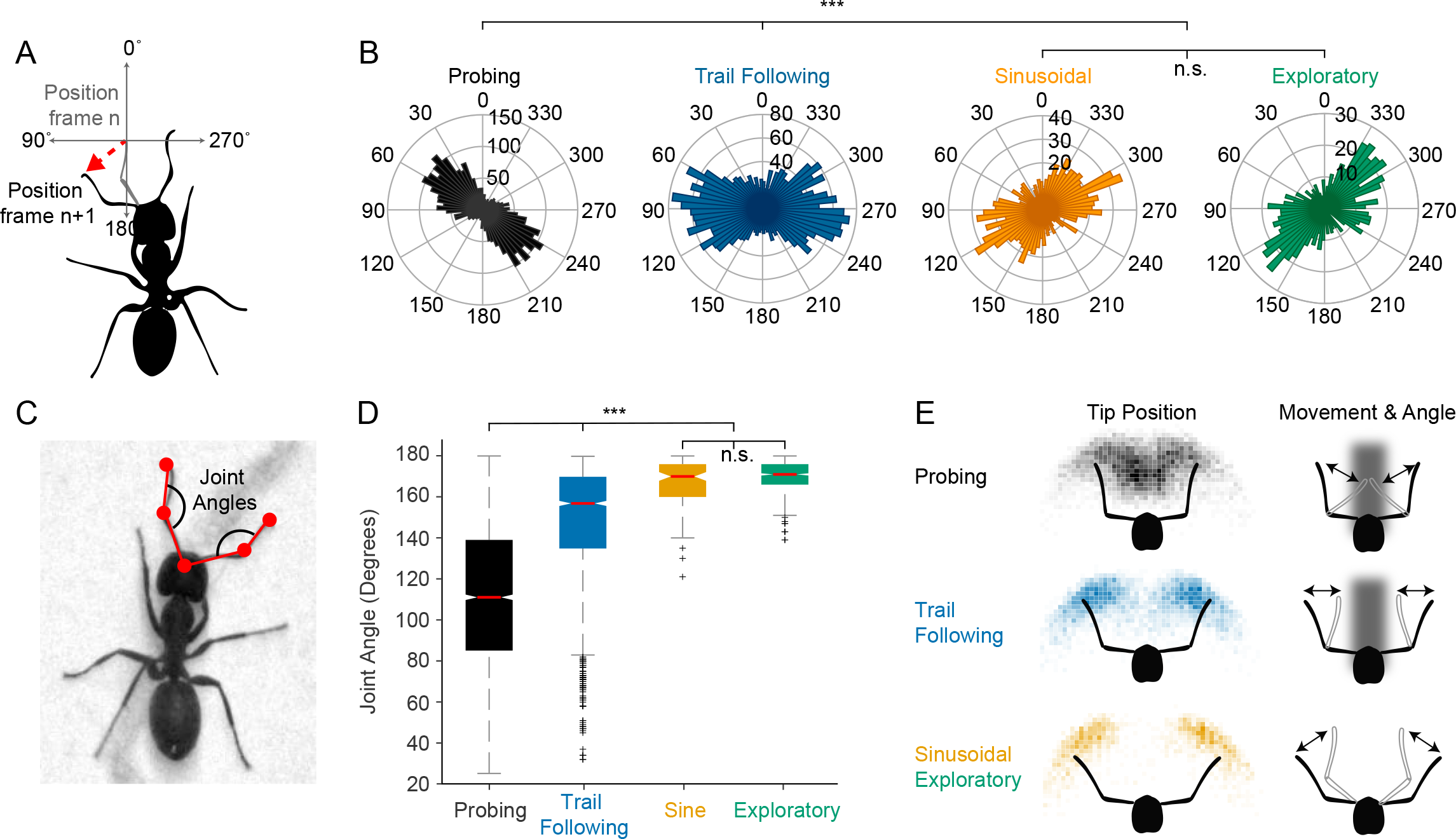
Pattern of antennae movement and joint angles differ during behavioral modules. A: Schematic showing the instantaneous angle of movement of the antenna tip between frames. B: Angle histograms of instantaneous movement angles for each of the behavioral modules (n=29 ant Trail Tracking; n=7 ant Exploratory; Probing: 3514 data points, Trail Following: 2998 data points, Sinusoidal: 1161 data points, Exploratory: 851 data points). All behaviors show distinct distributions of antennae movements except Sinusoidal and Exploratory (Kolmogorov-Smirnov, two-tailed, p < 0.0001 and p = 0.224 respectively, Bonferroni corrected, alpha= 0.008). *p < 0.05; **p < 0.01; ***p < 0.001. C: Schematic showing the points marking the head, pedicel, and antenna tip. Two lines between these three points were used to calculate the angle between the flagellar segment and the scape for both the left and right antennae. D: Joint angles during the trail tracking behavioral modules (n=7 ants; Probing: 4030 data points, Trail Following: 1922 data points, Sinusoidal: 138 data points) and Exploratory behavior (n=4 ants; 814 data points). All behavioral modules show distinct antennae joint angle distributions except for the comparison of Sinusoidal and Exploratory behaviors (Kolmogorov-Smirnov, two-tailed, all pairs p<0.0001 except Sinusoidal and Exploratory: p=0.009, Bonferroni corrected, alpha= 0.008). E: Summary of the movement angles, joint angles, and tip positions in each behavioral module.

The antennae contain an elbow-like joint between the scape and the flagellum, which changes its angle to help control the position of the antenna tip (Fig. 4C). We measured the distribution of joint angles in each of the behavioral modules to fully describe the movement of the antennae. Again, we see three broad groups (Fig. 4D, Kolmogorov-Smirnov, Bonferroni corrected, all pairs p<0.0001 except Sinusoidal and Exploratory: p=0.009). Probing displayed the widest range of angles (interquartile range: 139-85 degrees), indicating swinging of the flagellar segment inwards and outwards from the head. Exploratory and Sinusoidal behaviors have the smallest range of joint angles, concentrated at the largest angles (interquartile range: 176-160 degrees), indicating a rigid, outstretched antenna. Intermediate values of antennae angles were observed during Trail Following behavior (interquartile range: 170-135 degrees). Taken together, these data suggest a model of how the antennae move during these different behavioral modules, which is summarized in Fig. 4E. These findings show that behavioral modules classified with body movement parameters are associated with different strategies for antennae usage.

### During Trail Following, Ants Use Bilateral Information to Turn Towards Odor Signals

To study the bilateral use of antennae during tracking and to better understand how antenna signals affect the body position, we focused on the Trail Following behavioral module, when the ant tracks the trail closely and accurately. During Trail Following, the ant largely maintains the odor trail between its two antennae, as seen on the representative plot of antennae tip positions over time (Fig. 5A). To quantify the duration and degree of antennal tip contact with the trail, we measured the peaks and widths of trail overlap and compared it to those during Probing behavior (Fig. 5B). As shown, the Trail Following odor peak widths, which represent the amount of time an antenna tip spent on the trail, are much shorter than those during Probing (Fig. 5C, Wilcoxon Rank-Sum, p<0.0001). The peak heights, which represent the maximal extent of overlap between the trail and an antenna tip, are also higher during Probing behavior (Fig. 5D, Wilcoxon Rank-Sum, p<0.0001). The smaller and shorter peaks in trail overlap during Trail Following are consistent with the idea that the ants are briefly bringing their antennae to the edge of the trail and then moving them away.

**Figure 5.**
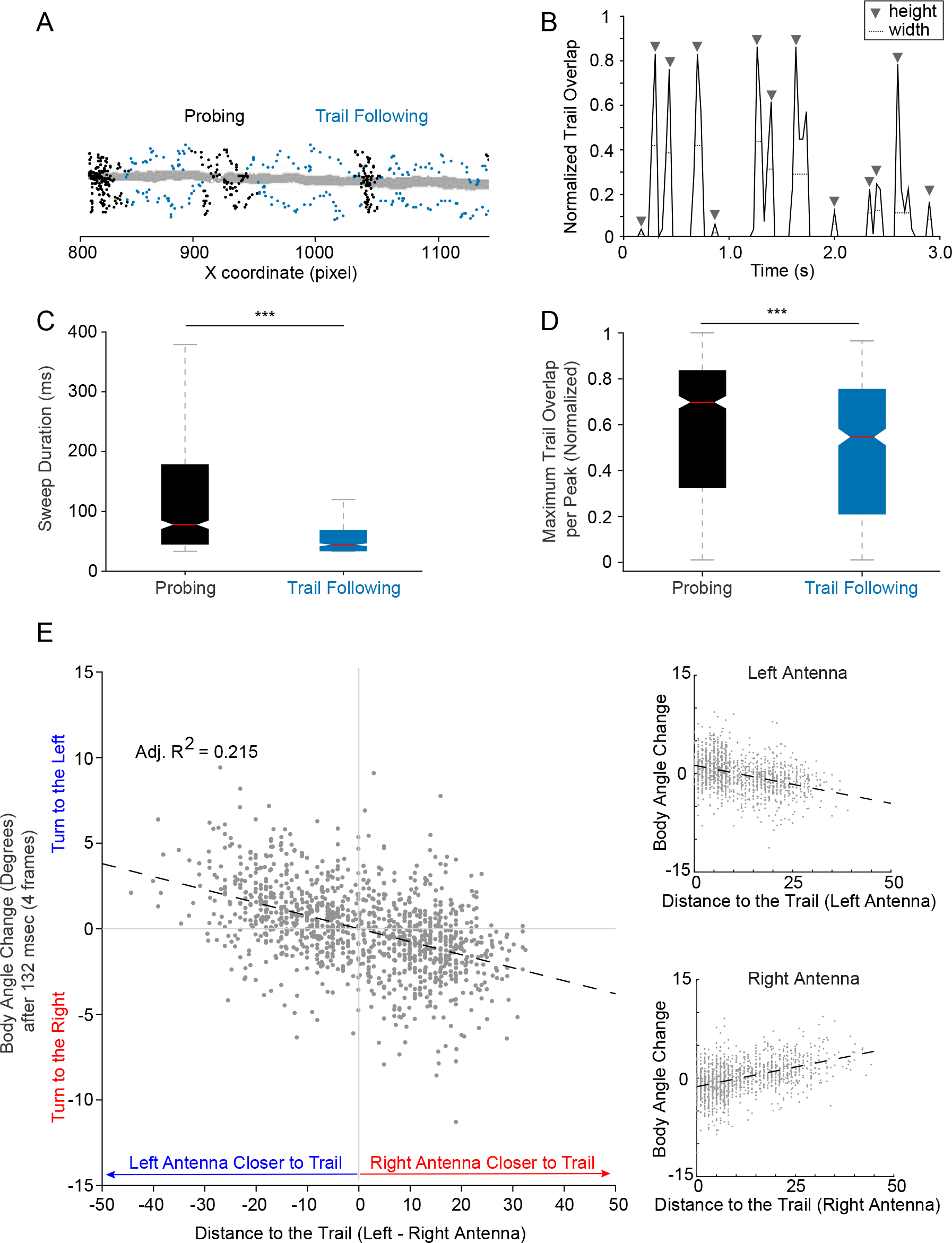
During Trail Following, ants track by turning towards small increases in odor concentration. A: Overlay of antennae tip positions during Probing (black) and Trail Following (light blue) with respect to the trail (grey) for a single ant trial. B: Normalized trail overlap for a single antenna over time showing peaks with widths and heights that reflect the extent of antenna tip overlap with the trail in time and space, respectively. Peak heights and widths of distinct encounters with the trail were marked computationally (grey triangles and dashed lines, respectively), see Methods for details. C. Quantification of peak widths for Probing and Trail Following (n= 29 ants; Probing: 863 peaks, Trail Following: 561 peaks). Peak widths in trail overlap were higher during Probing (median difference of 33.6 ms, Wilcoxon Rank-Sum, one-tailed, p < 0.0001). Outlier values were omitted from the Probing data plot to facilitate the comparison. *p < 0.05; **p < 0.01; ***p < 0.001. D: Quantification of peak heights for Probing and Trail Following (n=29 ants; Probing: 863 peaks, Trail Following: 561 peaks). Peak heights were larger during Probing (median difference of 0.15, Wilcoxon Rank-Sum, one-tailed, p < 0.0001). E: Linear regression on Trail Following behavior (1408 data points) showing the effect of antennae tip distances to the trail (left minus right) on turning angle (after 4 frames). When the left antenna is much closer to the trail, ants turn to the left and vice versa (Adjusted R2=0.215, F-test, p < 0.0001). Insets show the individual relationships between each antenna and turning angle (after 4 frames) (r = −0.39 and r = 0.44 for the left and right antenna, respectively).

We created a linear model to examine the relationship between odor sensation and changes in ant movement. Specifically, this model relates the antennae tip positions during Trail Following and the body angle change four frames later. This amount of delay (133 ms) was found to yield the best fit based on the adjusted R^2^ value. In this regression, the explanatory variables are the distances from both antenna tips to the trail separately, excluding frames where both antenna tips have zero overlap with the trail (*i.e.*, zero estimated odor concentration). Under these conditions, we obtained the relationship: ‘Angle Change’ = − 0.27 + 0.086*(‘Right Distance’) − 0.060*(‘Left Distance’), where both independent variables were necessary (t-test, p < 0.0001) and the model was significant (F-test, p < 0.0001, adjusted R^2^ = 0.215). The correlation between the body angle change and the left antenna distance to trail was r = −0.39 and the correlation of this angle with the right antenna was r = 0.44 (Fig. 5E insets). When plotting the differences between the left and right antennae tip distances, we see that the largest changes in angle occur when bilateral signaling is heavily biased towards one antenna (a large difference value). Based on the way we have defined the body angle, a positive change corresponds to a turn to the left. Thus, when the left antenna comes closer to the trail, the ant turns to the left, and a complementary relationship holds for the right antenna (Fig. 5E). This would result in a movement that keeps the trail centered between antennae.

### Ants show a difference in right/left antennae usage

The above data indicate that the two antennae are frequently at different distances from the odor trail, presumably conveying distinct odor concentrations. To determine if the movements of two antennae were coordinated, we calculated a correlation value between the distances of the two antennae from the trail for Probing and Trail Following, the only behavioral modules where the ant is close to the trail. Antennae positions were significantly anti-correlated in both behaviors (Fig. 6A and B, t-test, p<0.0001). The same result also holds when using the trail overlap variable instead of the distances of the antennae to the trail (<u>Movie 3,</u> Fig. S5). Furthermore, cross correlations between the left and right antennae distances to the trail at different frame lags reveal a periodic relationship between the two antennae (Fig. S5). This result reflects the periodic nature of the raw distance time series (Fig. 6A), which resembles two sine waves with different phases. To evaluate whether this anti-correlation exists in other behavioral modules, we repeated this analysis for distances of the two antennae from the body axis of the ant. Antennae tip positions relative to body axis were significantly anti-correlated during all behavioral modules except Trail Following (Fig. 6B, t-test, p<0.0001 except Sinusoidal: p=0.047 and Trail Following: p=0.096). Trail Following showed much lower anti-correlation values when using antennae tip to body axis distances compared to tip to trail distances. This suggests that, in Trail Following, alternating movements of the body axis shift the antennae tips towards and away from the trail (<u>Movie S1</u>). This is opposed to other behavioral modules, where the antennae themselves alternate about the body axis. Overall, across behavioral modules, movement of the two antennae is anti-correlated regardless of how and whether an ant is tracking an odor trail.

**Figure 6.**
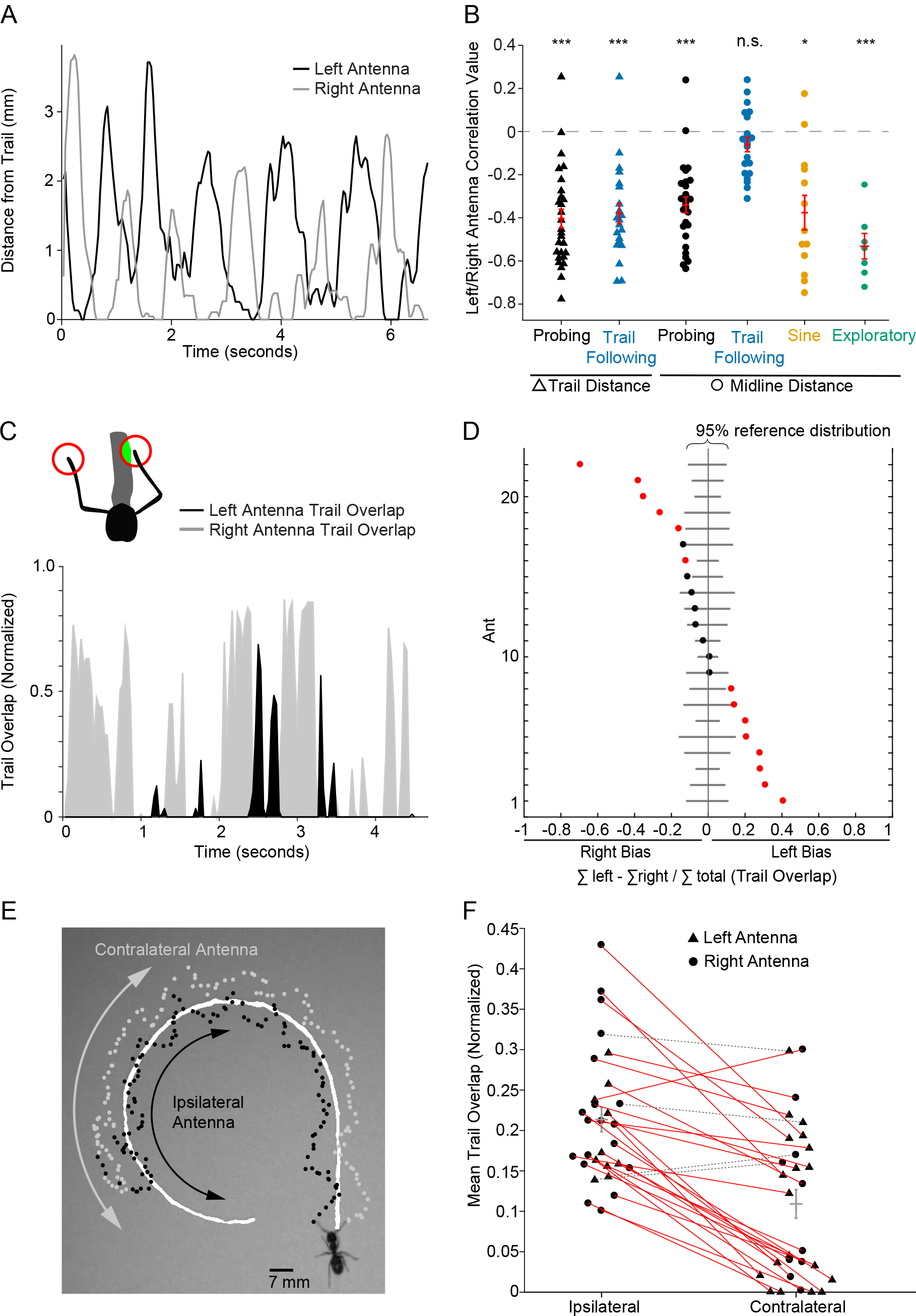
Antennae pairs show alternation and bias. A: Plot of right (grey) and left (black) antennae distances to the trail over time during Probing behavior for a single ant trial shows clear alternation. B: Correlation values for left and right antennae distance to the trail during Probing and Trail Following behaviors (triangles, t-test, two-tailed, p < 0.0001), and distance of the antennae to the ants’ body axis for all behavioral modules (circles, t-test, two-tailed, p < 0.0001 except Sinusoidal: p = 0.047 and Trail Following: p = 0.0964). Each point represents a behavioral ‘bout’ of at least 1 second (Probing bouts=28, Trail Following bouts=22, Sinusoidal bouts=13, Exploratory bouts=7). Statistical significance was evaluated using a t-test after Fisher’s z-transformation. Mean and SEM for each distribution is shown in red. *p < 0.05; **p < 0.01; ***p < 0.001. C: (Top) A diagram showing how trail overlap values were obtained. (Bottom) Plot of right (grey) and left (black) antennae trail overlap values over time for a single ant shows a clear bias towards the left antenna. D: A bias index was defined as the relative difference of total trail overlap values between the left and right antenna of a single ant after tracking a straight-line trail (n=22). The observed bias is shown as black and red dots (not significant and significant, respectively). Solid lines show the distribution of left-right bias from a randomization of the data from each ant (100,000 iterations per ant). 15 ants showed a dominant antenna (p<0.05, two-tailed, bootstrap distribution), with 7 and 8 ants showing a significant right and left bias, respectively. E: Representative image plotting right (here contralateral, grey) and left (here ipsilateral, black) antennae tip positions of an ant tracking a curved line trail.

Additionally, we observed that one antenna within a pair often overlaps significantly more with the odor trail than would be expected by chance (*i.e.*, random permutation of the same data, example raw data in Fig. 6C). We defined a bias index (*i.e.*, relative difference of trail overlap values, Fig. 6D) for each ant run. The population of ants analyzed showed a continuum of bias from very strongly left biased to very strongly right biased (Fig. 6D, 8 of 22 left-biased, 7 of 22 right-biased, p-value <0.05, bootstrap distribution). Since odor is encountered in discrete events when the antenna comes in contact with the trail, we also tested whether data were significantly different than expected if the total odor were randomly distributed in blocks equal to the number of peaks present in the trail overlap data (Fig. S6A, 9 of 22 biased, p-value <0.05, bootstrap distribution). In looking at the features of the peaks in the trail overlap data, it appears the bias arises from differences in the number of peaks, peak widths, and peak heights between the left and right antenna, but each of these factors contributes differently in those ants which show a bias (Fig. S6B).

This continuum of bias, found in straight line trail tracking ants, was not present in ants tracking trails of various degrees of curvature (from radius 3 cm to 6 cm, example in Fig. 6E). In curved trails, the biased antennae were nearly always ipsilateral to the inner curvature of the trail (Fig. 6F, 24 of 29 ants significantly ipsilaterally biased, 1 of 29 significantly contralaterally biased, Wilcoxon Sign-Rank, p < 0.05). These data suggest that there are biases in left-right usage among ants, but these biases may be context dependent.

Taken together, this shows that the left and right antennae are used in a non-redundant manner during tracking and that ants take advantage of bilateral symmetry during tracking behaviors to gather information about the environment.

### Single antenna ants show behavioral compensation through altered antenna usage

To test the importance of bilateral antennae information for tracking behavior, we removed a single antenna from ants and tested their ability to track trails (2-72 hours after surgery, n=6 ants, 14 trials). We compared the single antenna of these ants to the left antenna of our previously analyzed tracking ants. The single antenna ants showed a wider distribution of antennae tip positions, including positions farther away from the head and closer to the midline compared to the distribution of control ants (Fig. 7A and B, Kolmogorov-Smirnov Test, p < 0.0001). Furthermore, the single antennae often extended across the midline, which was much less apparent in control ants (Fig. 7A, red pixels). To quantify this difference, we compared the percent of time when an ant’s antenna crossed the midline and found single antennae crossed ~3-fold more often (Fig. 7C, Wilcoxon Rank-Sum, p=0.009). This raised the question of whether these different distributions impacted trail sampling. We focused this analysis on Trail Following, when the ant was closely tracking the trail, and quantified the percent of time an antenna extended beyond the contralateral side of the trail (Fig. 7C). The single antennae crossed the trail ~7-fold more often (Wilcoxon Rank-Sum, p=0.005). This shows that single antenna ants adapt to amputation by changing the pattern of usage of the remaining antenna to sample a larger physical space around themselves and around the trail.

**Figure 7.**
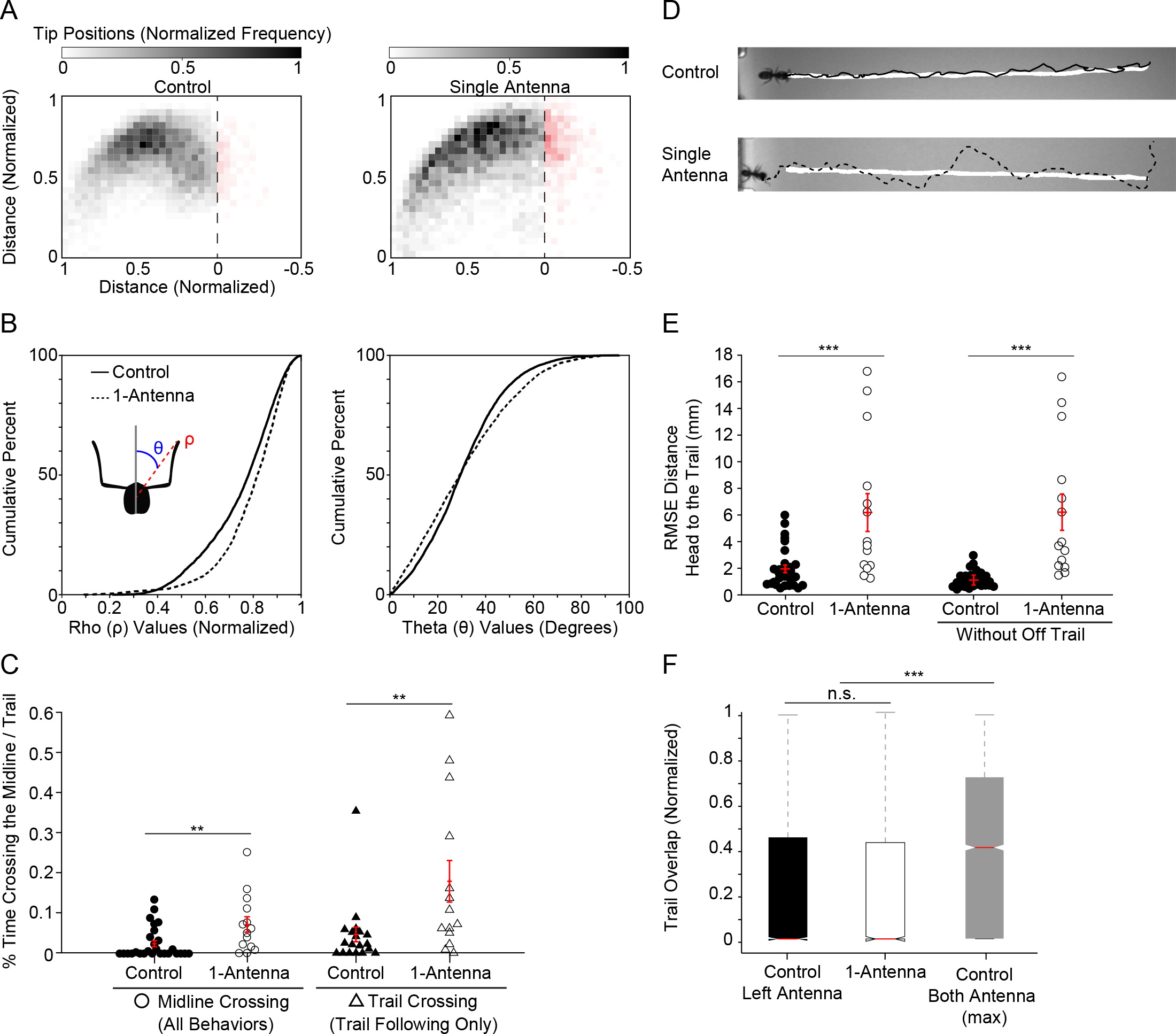
Single antenna removal results in reduced tracking accuracy and altered antenna usage. A: Normalized antenna tip positions from 29 control ants (left antenna only; 7945 data points) and 14 single antenna ants (6 ants over 14 trials; 4693 data points). Red pixels mark positions where the antenna tip crosses the body axis. B: Cumulative percent plots for control and single antenna ants of the distance of each antenna to the head (ρ; left) and the angle between the body axis and line connecting the head and the antenna point (θ; right). For both variables, the two distributions were statistically distinct (Kolmogorov-Smirnov, two-tailed, p < 0.0001). C: Percent of time when an ant’s antenna extended to the contralateral side of the body axis for control ants (n = 29, filled circles) and single antenna ants (n=14, open circles). Percent of time during Trail Following when an ant’s antenna extended beyond the contralateral edge of the trail for control ants (n = 19, filled triangles) and single antenna ants (n=14, open triangles); only ants with at least 30 frames of Trail Following behavior were considered. In both cases, single antenna ants crossed their body axis and the trail with their antenna more often than control ants (2.9-fold more and 6.8-fold more, respectively; Wilcoxon Rank-Sum, one-tailed, p < 0.01). Mean and SEM for each group is shown in red. *p < 0.05; **p < 0.01; ***p < 0.001. D: Representative images plotting ant head position over time for a control (solid line) and single antenna (dashed line) ant during trail tracking (trail pixels are labeled white). E: Comparison of the root mean square error (RMSE) of the ant head to trail distance for control (left antenna, n=29) and single antenna (n=14) ants. The same comparison is shown with data excluding Off Trail behavior from both groups. In both cases, the single antenna ants have a higher RMSE than control ants (Wilcoxon Rank-Sum, one-tailed, p < 0.0001). Mean and SEM for each group is shown in red. F: Trail overlap for the control ants (left antenna only), single antenna ants, and control ants (maximum of left and right antenna values per frame). The control ants’ left antennae and the single antennae show similar overlap with the trail (Wilcoxon Rank-Sum, one-tailed, p = 0.685), but the control ants’ maximal overlap is significantly larger (Wilcoxon Rank-Sum, one-tailed, p < 0.0001).

Next, we evaluated how accurately the single antenna ants were able to follow the odor trail by plotting the distance of the head to the trail over time. We compared the root mean square error (RMSE) of this distance and found that single antenna ants had significantly higher RMSE (Fig. 7D and E, Wilcoxon Rank-Sum, p < 0.0001). The side on which errors occurred was not consistently correlated with the position of the removed antenna (data not shown).

Finally, we compared the trail overlap from the single antenna and control ants (Fig. 7F, left and middle boxplots). Surprisingly, we found these estimated odor distributions to be similar (Wilcoxon Rank-Sum, p = 0.685), suggesting that the wider sampling area of single antennae help to compensate for larger head to trail deviation. However, as shown previously, single antenna ants track less accurately (Fig. 7E); given that the two antennae are used non-redundantly (Fig. 6), control ants receive much more odor information over time. Therefore, we selected, from control ants, the larger of the two antennae trail overlap values at all time points as a comparison (Fig. 7F, right boxplot). Indeed, single antenna ants show far less overlap with the trail compared to this bilateral signal (Wilcoxon Rank-Sum, p < 0.0001). Therefore, having two antennae allows the ant to be in contact with the trail more continuously.

## Discussion

In this study, we quantified and modeled the strategies of tracking behavior and antennae usage in ants (*Camponotus pennsylvanicus*). We have shown, for the first time, that ants use a variety of behavioral strategies during trail tracking, each of which has a distinct and stereotyped pattern of antennae sampling. In all trail tracking behaviors, we found that the left and right antenna play non-redundant roles in odor detection: they generally move in anti-correlated fashion and ants often exhibit a bias in which antenna is closer to the odor trail. This observation suggests lateral specialization, and moreover, the use of complex strategies for odor detection. We further examined the influence of bilateral comparisons over an extended trail tracking episode, and we found that ants show signatures of edge detection and use information from both antennae to guide turning and accurately follow the trail. Removal of one antenna impairs tracking but also results in adaptation of antenna sampling, suggesting ants can adopt motor strategies to partially compensate for antenna loss.

### Antennae show distinct patterns of movement and placement during odor tracking

Quantitative studies have led to a greater understanding of olfactory behavior in many species (*e.g.*, Gomez-Martin, 2011; Riffell, 2014; Khan, 2012). Although chemical communication and trail following in ants have been studied for many decades, there is very little known about sensorimotor strategies at the level of antenna. Ants have relatively long sensors that are highly motile. As a first step in being able to model their olfactory behavior, we looked at antennae movement during trail tracking at high spatial (0.22 mm/pixel) and temporal resolution (30 fps). We found that antennae are highly dynamic whether tracking a trail or exploring (Fig. 1C). However, during tracking, antennae sample a much larger area of space (Fig. 1B). Additionally, by measuring ant speed and body movement relative to the trail, we were able to classify behavior into three discrete components (or behavioral modules, Fig. 2), similar to classifications done in other invertebrates (Green, 1983). These three behavioral modules showed distinct patterns of antennae sampling.

The first behavioral module, *Sinusoidal*, was characterized by periodic movement along the odor trail. This type of movement has been observed in ants (Holldobler and Wilson, 1990) as well as a range of species (Martin, 1965; Borst and Heisenberg, 1982; Gomez-Martin, 2010; Khan, 2012; Porter, 2007). It is thought to represent a search pattern along an odor trail or gradient, where temporal odor information is used to stay centered on a signal. In our analysis, we found that the antennae tip positions and movements in this behavioral module were restricted to a smaller area (when compared to Probing or Trail Following) and were indistinguishable from antennae positioning during Exploratory behavior (Fig. 3B). Therefore, odor sampling during Sinusoidal behavior is accomplished by alternating movements of the ant itself along the trail while keeping its antennae outstretched.

The second behavioral module, *Probing*, described ants that were stationary and within sampling distance of the trail. During Probing, the ant’s antennae had the largest area of sampling (Fig. 3B) and were most densely focused on the trail itself (Fig. 5A). Additionally, ants swept their antennae back and forth towards the head (Fig. 4B). These observations are consistent with an intensive sampling of odor to examine and identify its relevance. This behavioral module is not exclusive to encountering the experimenter-drawn trail; ants not given a trail would infrequently engage in a Probing-like behavior as well (data not shown).

The third behavioral module, *Trail Following*, was characterized by an ant following the trail with high accuracy. The antennae tip distribution largely excluded an area at the midline that was approximately the width of the trail (Fig. 3A). Antennae moved in a direction perpendicular to the trail (Fig. 4B), and the short duration and extent of antennae tip trail overlap suggest this behavior involves edge detection (Fig. 5C and D). Evidence of edge detection during tracking has been seen in other animals such as bats and rats (Yovel, 2010; Khan, 2012).

In these three behavioral modules, the antennae tips displayed distinct spatial distributions, and as such they would gather and convey to the brain different information about trail odor (Fig. 3D). Previous studies in various species have analyzed large-scale antennal movements in response to stimuli (*e.g.* Huston, 2015; Pribbenow, 1996), but finer movements and distinct strategies are largely uncharacterized. Given the importance of understanding how sensors are moved to sample the environment and gather information to guide behavior, our models in ants provide an important basis for future studies.

### Ants use bilateral information to guide turning during Trail Following

Previous experimental and theoretical works have proposed a model of tropotaxis to explain how insects track odor trails (Martin, 1965; Hangartner, 1967; Borst and Heisenberg, 1982; Calenbuhr and Deneubourg, 1992; Louis, 2008). In Argentine ants, behavior at trail bifurcation points could be predicted based on estimated concentration difference between antennae (Perna, 2012). We tested this idea in Trail Following behavior by using the distance of the trail to the antenna as a proxy for odor signal. In a linear model, we found that the direction and magnitude of the turning angle of the ant could be partially explained by the difference in trail distance between antennae (Fig. 5D, Adjusted R^2^=0.215). We found that larger differences in estimated odor between the two antennae led to larger turning angles towards the trail (Fig. 5C). The best model used a 133 ms (4 frame) delay between odor sensation and subsequent turns. This latency for motor implementation is similar in timescale to adult drosophila, which have been shown to redirect flight heading within 250 ms of detecting an odor plume (Burdick and Dickinson, 2006). These results demonstrate that ants use bilateral differences in antenna signals to track odors during Trail Following behavior. We note, however, that putative antennae concentration difference can only partially predict ant movement, and many other factors are likely to be involved in motor decision making.

### Antennae have non-redundant roles in sampling odor trails

Given the strong evidence for the importance of bilateral comparisons, we asked whether each antenna within a pair is used differently during trail tracking. We looked at the correlation between antennae using either their distance to the trail or to the ant’s body axis. In all behavioral modules, the antennae exhibit a strong anti-correlation (Fig. 6B). Assuming odor information is typically highly variable in time and space, keeping the two odor sensors in different locations would increase the efficiency of detecting odors and sampling odor space. As far as we know, this is the first evidence of stereotyped coordination between antennae during odor tracking. Except for Trail Following, these anti-correlations arise from alternating movements of the antennae with respect to the body axis. In Trail Following, antennae predominantly show anti-correlation with respect to the trail but not to the body axis. One possibility is that the antennae movement driving the anti-correlation with respect to the trail comes from changes in the angle of the body axis rather than movements of antennae relative to the body axis. Indeed, careful inspection of the videos shows numerous examples of the ant making small adjustments in body angle with respect to the trail during Trail Following (<u>Movie S1</u>). Thus, like Sinusoidal behavior, the body movement of the ant is an important variable involved in positioning the antennal sensors of odor.

In addition to the anti-correlation between antennae, we found a continuum of antennae bias in trail sampling. In our study, we found a large fraction of ants (15 out of 22) where one antenna overlapped significantly more with the trail than the other antenna during a tracking run (Fig. 6C). In some ants, the left antenna was strongly biased (8 of 15) and in others the right was strongly biased (7 of 15), similar to studies showing a continuum of handedness in turning bias in flies (Buchanan, 2015). This bias was radically altered when the odor trails were curved as opposed to straight (Fig. 6E and F). When following curved trails (either to the right or the left), the bias was strongly towards the antenna ipsilateral to the inner curvature. This result could be interpreted in two ways. First, any inherent bias in sampling *(i.e*., ‘handedness’) can be masked by trail features that impose a greater bias in antennae overlap. Or, second, the pattern of antennae sampling used by ants is not inherent but rather context dependent, such that ants use different sampling strategies in different situations. Regardless, the presence of a bias suggests lateral specialization within an antennae pair that affects odor sampling on simple trails. This fits with evidence from a variety of studies which has shown a lateralization of insect brains and behaviors (Buchanan, 2015; Wes, 2001; Letzkus, 2006; Rogers, 2013; Frasnelli, 2012).

### Single antenna removal mildly impairs tracking and results in changes in antenna position and sampling

Qualitative and quantitative studies have repeatedly shown that single antennectomy results in a modest decrease in tracking accuracy in insects (Martin 1965; Hangartner, 1967; Borst and Heisenberg, 1982; Duistermars, 2009; Steck, 2010; c.f., Lockey and Willis, 2015). We found similar results: deviation from the trail (as measured by RMSE) was higher for single antenna ants (Fig. 7C). Ants were, however, still able to track trails along their entire length (15 cm), as has been seen previously, and which is thought to be due to angle corrections after successful temporal comparisons (klinotaxis). Interestingly, we saw altered antennae tip positions and sampling in ants after surgery (Fig. 7A and B). After 2-72 hours, the single antenna moved over a larger area of space, including extending across the ant’s midline and the contralateral edge of the trail, which is rarely seen in controls (Fig. 7B). This is consistent with evidence from drosophila that show adaptation within days after amputation (Wosnitza, 2013; Isakov, 2016; Muijres, 2017), as well as recent studies on ants showing robust performance across a range of olfactory behaviors after unilateral antenna amputation (Waxman, 2017). This resulting behavioral change in antennae movement may partially explain how the single antenna ant is able to maintain a level of estimated odor information similar to what one antenna of a control ant receives despite greater deviation from the trail (Fig. 7D).

## Conclusion

Here we have developed a better understanding of the multiple behavioral strategies ants use to track odor trails. This includes three distinct and stereotyped patterns of body and antenna movement and usage, which affect the amount of odor information received over time by each antenna and the ant brain. These descriptions provide insight into the general strategies ants use for trail tracking as well as evidence for how the two, distinct antennae move with respect to each other and relate to motor output. In future studies, behavioral analysis could be coupled to neural recordings to determine how odor inputs and proprioceptive information are transformed into motor outputs in simple, as well as more complex, odor trails.

## Acknowledgements

We thank all the members of the Murthy lab for their helpful discussions on experimental design and data analysis. We thank the members of the Traniello lab, especially Mario Muscedere and Fran Kamhi, for teaching us to collect, house, and dissect ants. We thank Ben de Bivort for his comments on the analysis, and Rob Johnson for his critical reading of the manuscript.

## Data and Software Availability

All data and software are available from the corresponding author, Ryan W. Draft, upon request.

## Movies

https://www.dropbox.com/sh/rl1f9pw1il1sx8s/AACkCVrHI_MXW-7VCoyAN_CKa?dl=0

### Movie 1. Antennae tip detection and marking

A tracking trial (631 frames) was background subtracted (green), cropped, and aligned. A search window was defined in the area above the head centroid and limited to pixels with values greater than a dynamic threshold (purple). Within the search area, the longest paths from the head point were used to find the antenna tips (crosses). Data were checked for errors and manually corrected.

### Movie 2. Parameterized ant tracking a 15-cm odor trail (30 fps, 0.22 mm/px)

An ant exits the holding chamber (left) and tracks an odor trail. Ant centroid (cyan dot), head point (magenta dot), antennae tips (blue circle, left tip; red circle, right tip), and body axis (white) were overlaid on the video. The boundaries of the trail threshold are shown in green.

### Movie 3. ‘Trail overlap’ during tracking

A tracking trial was cropped and aligned. Blue (left) and red (right) circles display the approximate area used for ‘trail overlap’ measurement. Bar plots show the ‘trail overlap’ values per frame, and a line plot of these values over time is shown above.

## Supplementary Movies

### Movie S1. Changes in body axis angle during Trail Following

Video clips (inverted for clarity) showing four ants during Trail Following behavior. Periodic changes in the body axis towards and away from the trail can be seen. Body angles were plotted and color-mapped to the maximum (red) and minimum (blue) angles during each bout.

---

**Figure S1.**
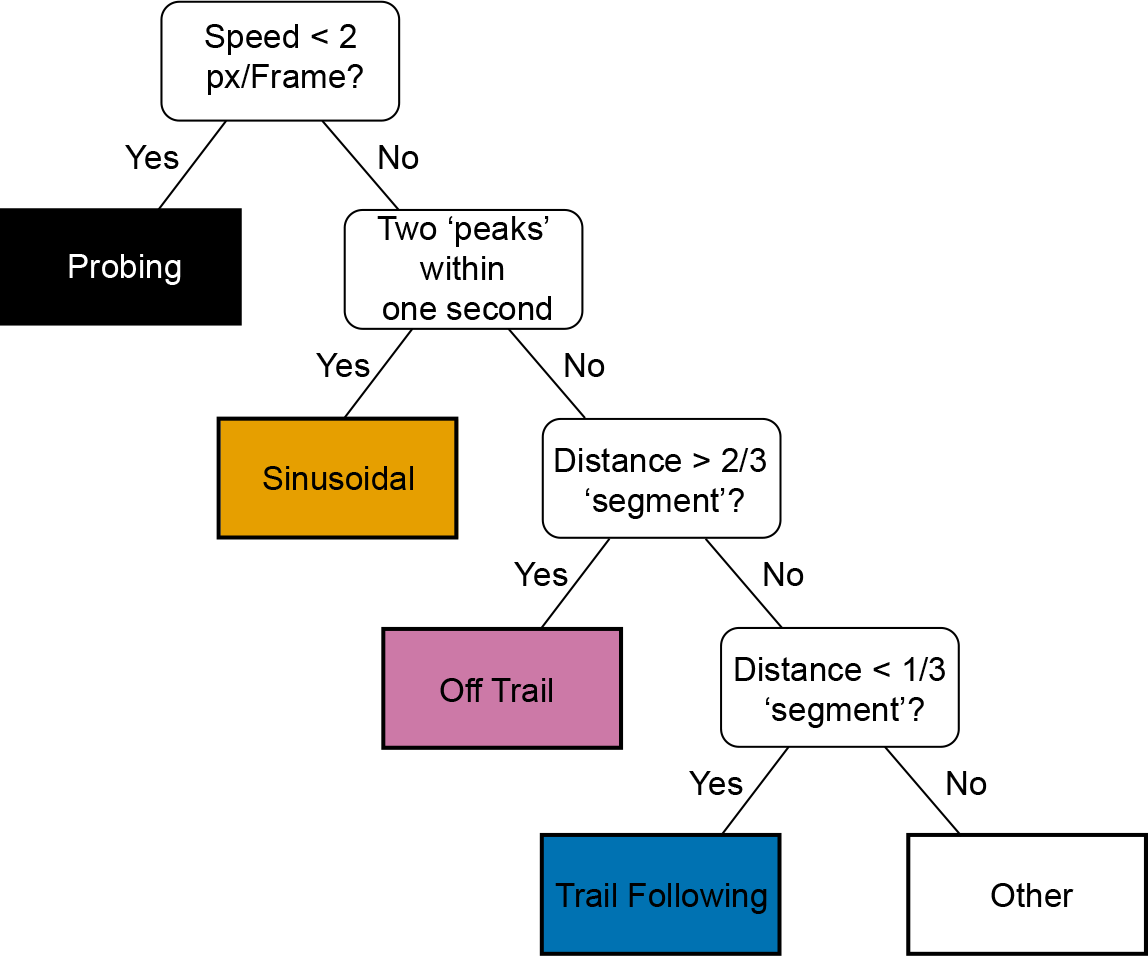
Major steps in the behavioral module classification algorithm. Flow chart showing the order and major criteria for behavioral categorization. First, Probing behavior was marked (speed < 2 pixels per frame for at least 5 frames). Second, Sinusoidal behavior was marked (two head-to-trail distance peaks within one second). Third, Off Trail was marked (head-to-trail distance greater than two-thirds of a body size based unit, or segment). Fourth, we identified Trail Following (head-to-trail distance less than one-third of a segment for at least 5 frames). Any remaining unclassified frames were marked as Other.

**Figure S2.**
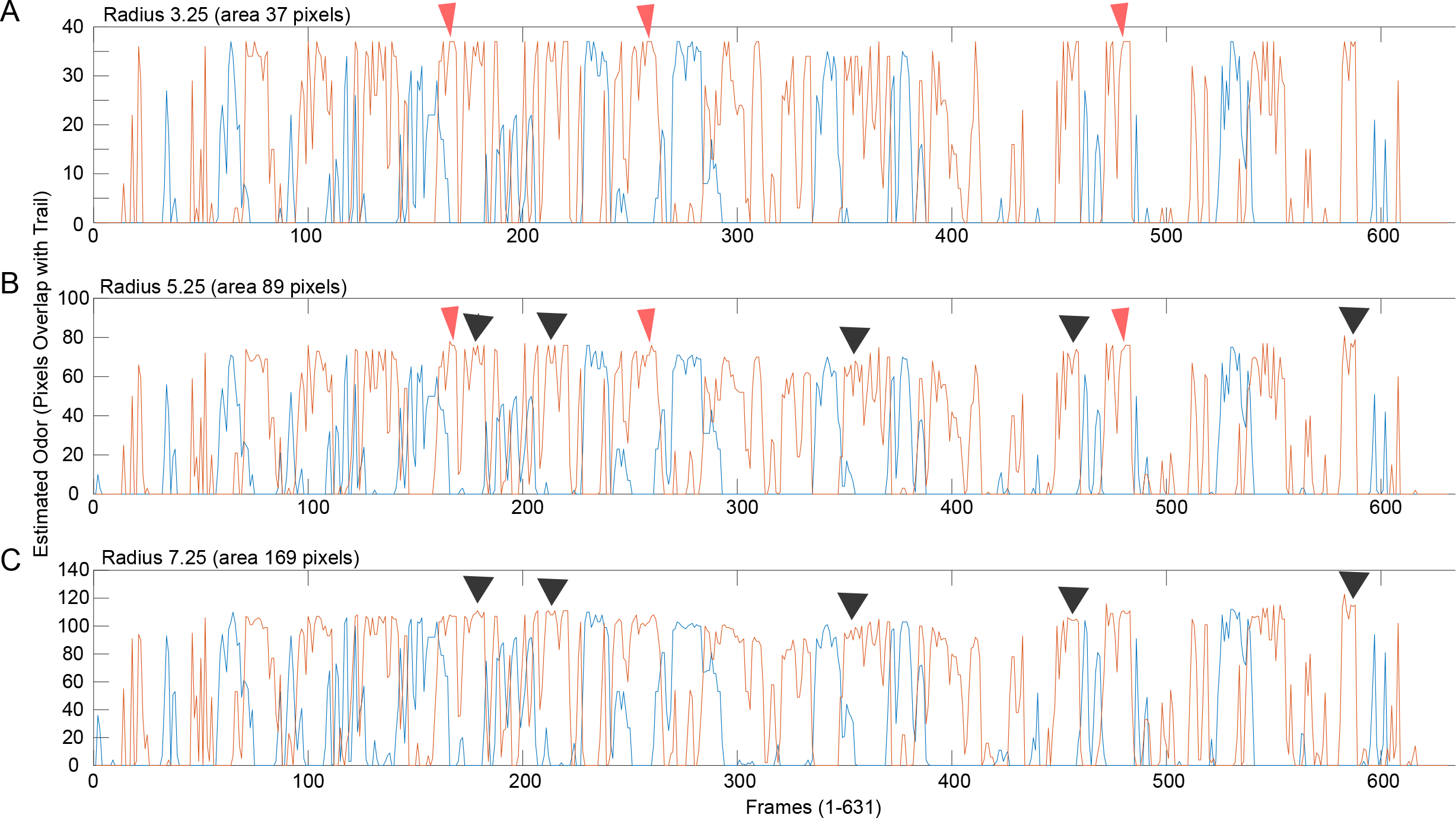
Effect of radius size on ‘trail overlap’ values. For a given ant trial (631 frames), we plotted ‘trail overlap’ over time for circles of radius 3.25 (top), 5.25 (middle), and 7.25 (bottom). Minor differences in the shape of the curves are apparent (right antenna: orange line, left antenna: blue line). Black arrowheads mark differences (information loss) occurring at the larger radius. Crimson arrowheads mark differences (information loss) occurring at the smaller radius.

**Figure S3.**
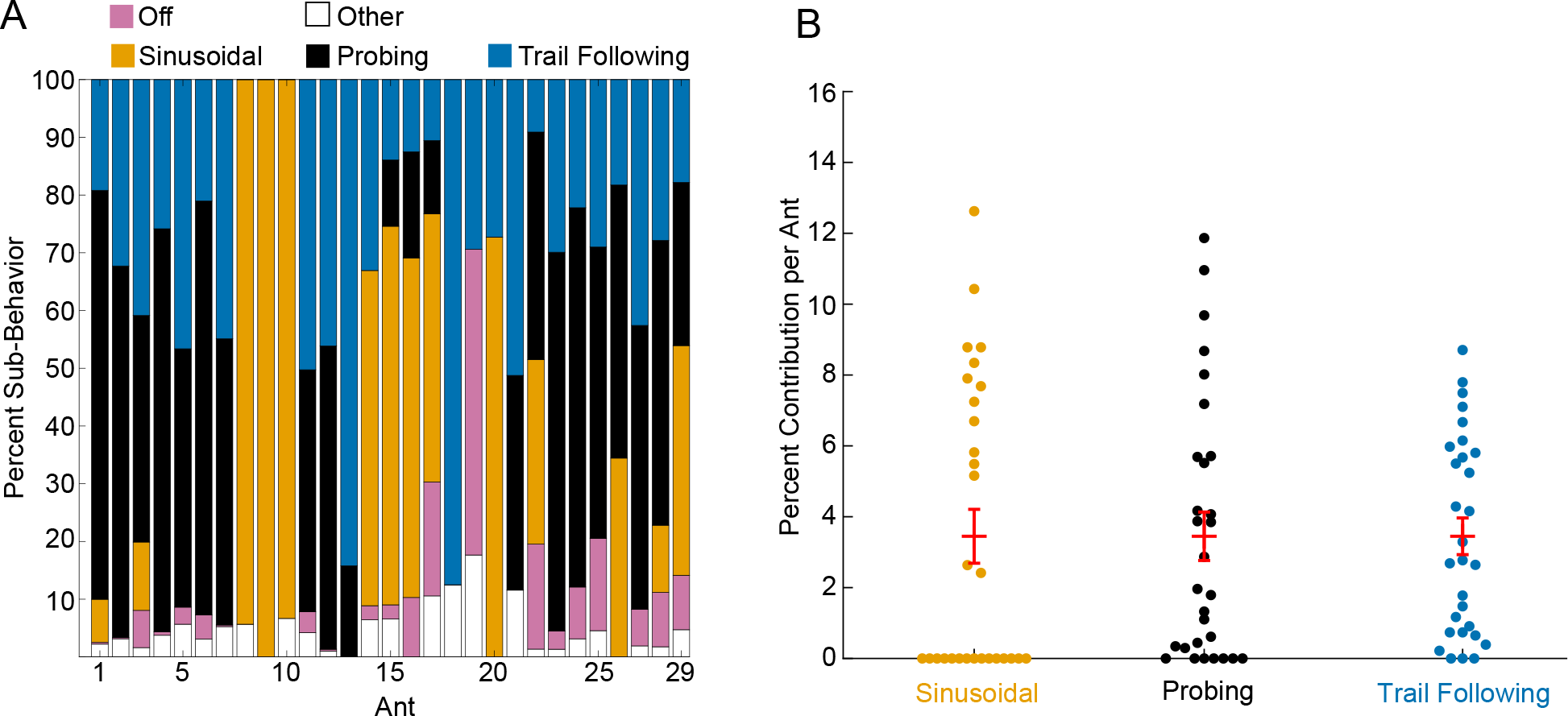
Percentage of behavioral modules per ant trial. A: Bar plot showing the percent of frames in a given trail-tracking trial that the ant spent in each behavioral module (Probing, Sinusoidal, Trail Following, Off Trail, and Other). Most ants (24 of 29) exhibited two or more of the specific trail tracking behavioral modules (Sinusoidal, Probing, or Trail Following). B: Plot showing the percent of frames each trail tracking ant (n=29) contributed to the total data analyzed for each behavioral module. As shown, each ant contributes a fraction of the data (mean 3.5 percent). Mean and SEM for each group is shown in red.

**Figure S4.**
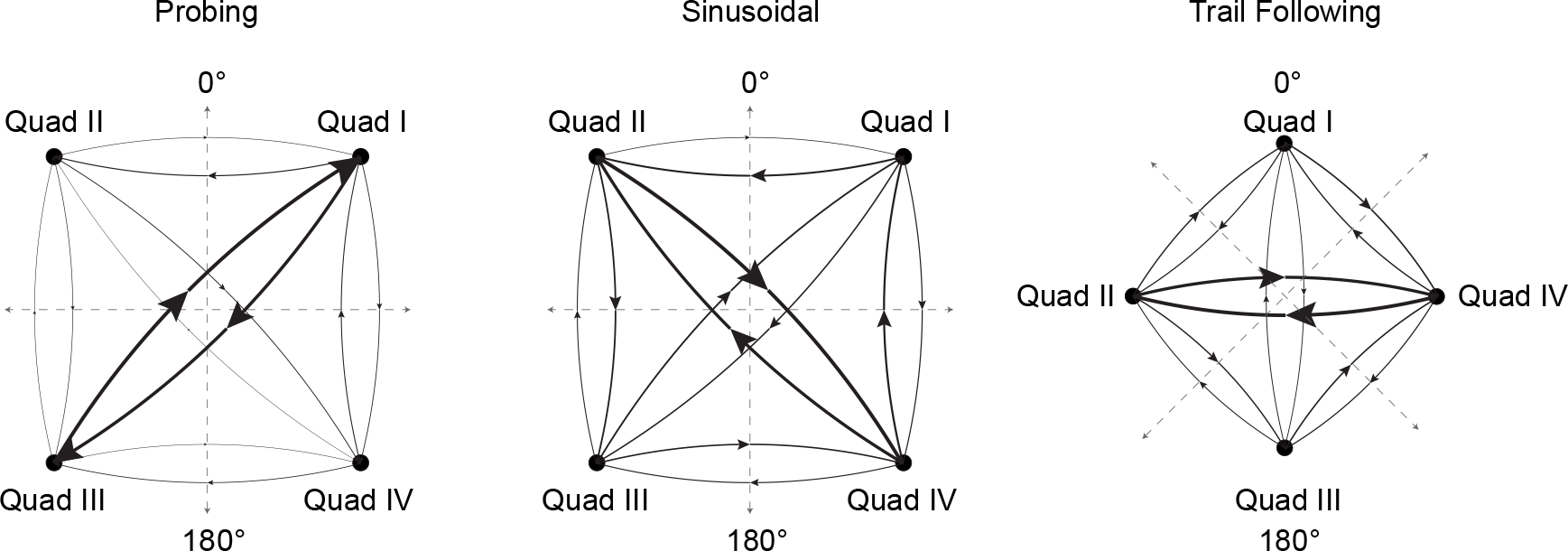
Connectivity plots for each behavioral module. Vectors of antennae tip movement from Fig. 4A were categorized based on the coordinate plane quadrant in which they reside. To evaluate how ants move their antennae over time, we quantified the transitions between these antennae movements binned by quadrant. The distribution of these transitions in movement is shown as relative line thickness in a connectivity diagram. The plots are dominated by a back and forth motion in two main directions for each behavior. Note that for Trail Following, axes were rotated by 45 degrees before binning the movement vectors to highlight connectivity between movements at 90 and 270 degrees (quadrants II and IV).

**Figure S5.**
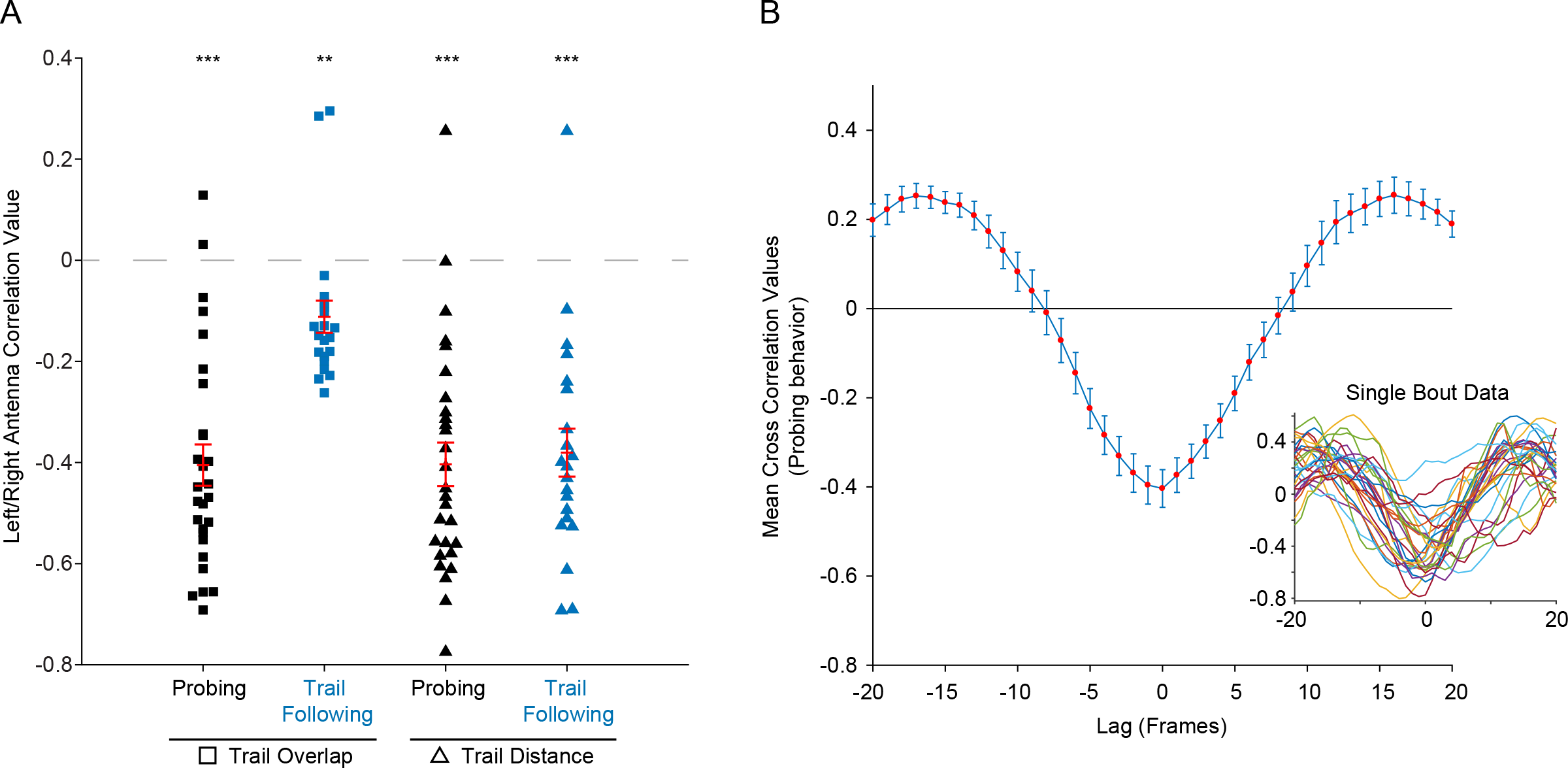
Cross correlations between antennae. A: Correlation values for left and right antennae trail overlap during Probing and Trail Following behaviors (squares, t-test, two-tailed, p < 0.0001 and p=0.0025, respectively), and distance of the antennae to the trail for the same behaviors (triangles, t-test, two-tailed, p < 0.0001). Each point represents a behavioral ‘bout’ of at least 1 second (Probing bouts=28, Trail Following bouts=22). Statistical significance was evaluated using a t-test after Fisher’s z-transformation. Mean and SEM for each distribution is shown in red. *p < 0.05; **p < 0.01; ***p < 0.001. B: Cross correlation comparing the left and right antennae distance to the trail during Probing for lags of +/− 20 frames. The inset shows the cross correlation results for each behavioral bout (n=28) and the large plot shows the mean correlation at each lag. Error bars represent SEM.

**Figure S6.**
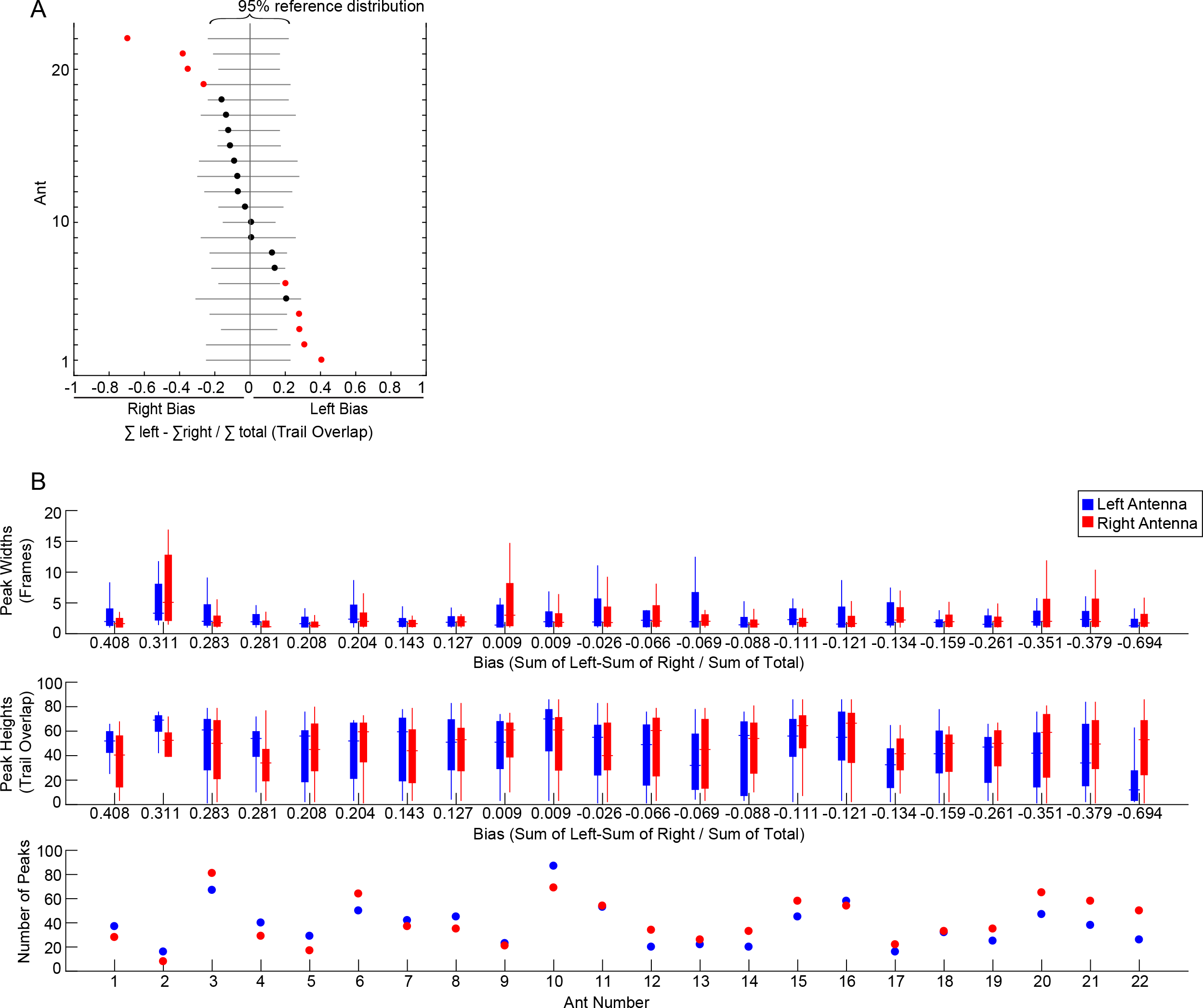
Analysis of left-right bias among antenna pairs. A: A bias index was defined as the relative difference of total trail overlap values between the left and right antenna of a single ant after tracking a straight-line trail (n=22 ants). Solid grey lines show the expected distribution of left-right bias for each ant that results from randomly distributing the sum of all trail overlap values (left and right) in blocks equal to the total number of discrete peaks in the ant trial (see Methods; 100,000 iterations per ant). The observed bias is shown as black and red dots (not significant and significant, respectively). 9 ants showed a dominant antenna (p<0.05, two-tailed, bootstrap distribution), with 5 and 4 ants showing a significant left and right bias, respectively. B: The peak widths (top), heights (middle) and number (bottom) from left (blue) and right (red) antennae trail overlap data (n= 22 ants). Ants differ in how each of these variables contribute to the total trail overlap values of the left and right antennae.

